# Increased Potency and Breadth of SARS-CoV-2 Neutralizing Antibodies After a Third mRNA Vaccine Dose

**DOI:** 10.1101/2022.02.14.480394

**Authors:** Frauke Muecksch, Zijun Wang, Alice Cho, Christian Gaebler, Tarek Ben Tanfous, Justin DaSilva, Eva Bednarski, Victor Ramos, Shuai Zong, Brianna Johnson, Raphael Raspe, Dennis Schaefer-Babajew, Irina Shimeliovich, Mridushi Daga, Kai-Hui Yao, Fabian Schmidt, Katrina G. Millard, Martina Turroja, Mila Jankovic, Thiago Y. Oliveria, Anna Gazumyan, Marina Caskey, Theodora Hatziioannou, Paul D. Bieniasz, Michel C. Nussenzweig

**Author notes:** equal contribution (Frauke Muecksch, Zijun Wang, Alice Cho). Address correspondence to: Theodora Hatziioannou; Paul D. Bieniasz; or Michel C. Nussenzweig.

## Abstract

The omicron variant of SARS-CoV-2 infected very large numbers of SARS-CoV-2 vaccinated and convalescent individuals^1–3^. The penetrance of this variant in the antigen experienced human population can be explained in part by the relatively low levels of plasma neutralizing activity against Omicron in people who were infected or vaccinated with the original Wuhan-Hu-1 strain^4–7^. The 3^rd^ mRNA vaccine dose produces an initial increase in circulating anti-Omicron neutralizing antibodies, but titers remain 10-20-fold lower than against Wuhan-Hu-1 and are, in many cases, insufficient to prevent infection^7^. Despite the reduced protection from infection, individuals that received 3 doses of an mRNA vaccine were highly protected from the more serious consequences of infection^8^. Here we examine the memory B cell repertoire in a longitudinal cohort of individuals receiving 3 mRNA vaccine doses^9,10^. We find that the 3^rd^ dose is accompanied by an increase in, and evolution of, anti-receptor binding domain specific memory B cells. The increase is due to expansion of memory B cell clones that were present after the 2^nd^ vaccine dose as well as the emergence of new clones. The antibodies encoded by these cells showed significantly increased potency and breadth when compared to antibodies obtained after the 2^nd^ vaccine dose. Notably, the increase in potency was especially evident among newly developing clones of memory cells that differed from the persisting clones in targeting more conserved regions of the RBD. Overall, more than 50% of the analyzed neutralizing antibodies in the memory compartment obtained from individuals receiving a 3^rd^ mRNA vaccine dose neutralized Omicron. Thus, individuals receiving 3 doses of an mRNA vaccine encoding Wuhan-Hu-1, have a diverse memory B cell repertoire that can respond rapidly and produce antibodies capable of clearing even diversified variants such as Omicron. These data help explain why a 3^rd^ dose of an mRNA vaccine that was not specifically designed to protect against variants is effective against variant-induced serious disease.

## Main

We studied the immune responses to SARS-CoV-2 mRNA vaccination in a longitudinal cohort of 43 volunteers with no prior history of SARS-CoV-2 infection^9,10^, who were recruited between January 21, 2021, and December 14, 2021, for sequential blood donation. Volunteers received either the Moderna (mRNA-1273; n=8) or Pfizer-BioNTech (BNT162b2; n=35) mRNA vaccines. The volunteers ranged in age from 23-78 years old 53% were male and 47% female (for details see Methods and Supplementary Table 1). Samples were obtained at the following time points: 1) 2.5 weeks after the prime; 2) 1.3 and 5 months after the 2^nd^ vaccine dose; 3) 1 month after the 3^rd^ dose.

### Plasma binding and neutralization

Plasma IgM, IgG and IgA responses to SARS-CoV-2 RBD were measured by enzyme-linked immunosorbent assay (ELISA)^9,10^. After a significant decrease in antibody reactivity during the 5 months following the second vaccine dose, anti-RBD IgG titers were significantly increased following a 3^rd^ dose of an mRNA vaccine (p<0.0001, Fig. 1a and Supplementary Table 1). The resulting titers were similar to those found 1.3 months after the 2^nd^ dose (p>0.99, Fig. 1a). IgM and IgA titers were lower than IgG titers and while IgM titers were unchanged during the observation period, IgA titers also increased significantly following a 3^rd^ vaccine dose (Extended data Fig. 1 and Supplementary Table 1).

**Fig. 1:**
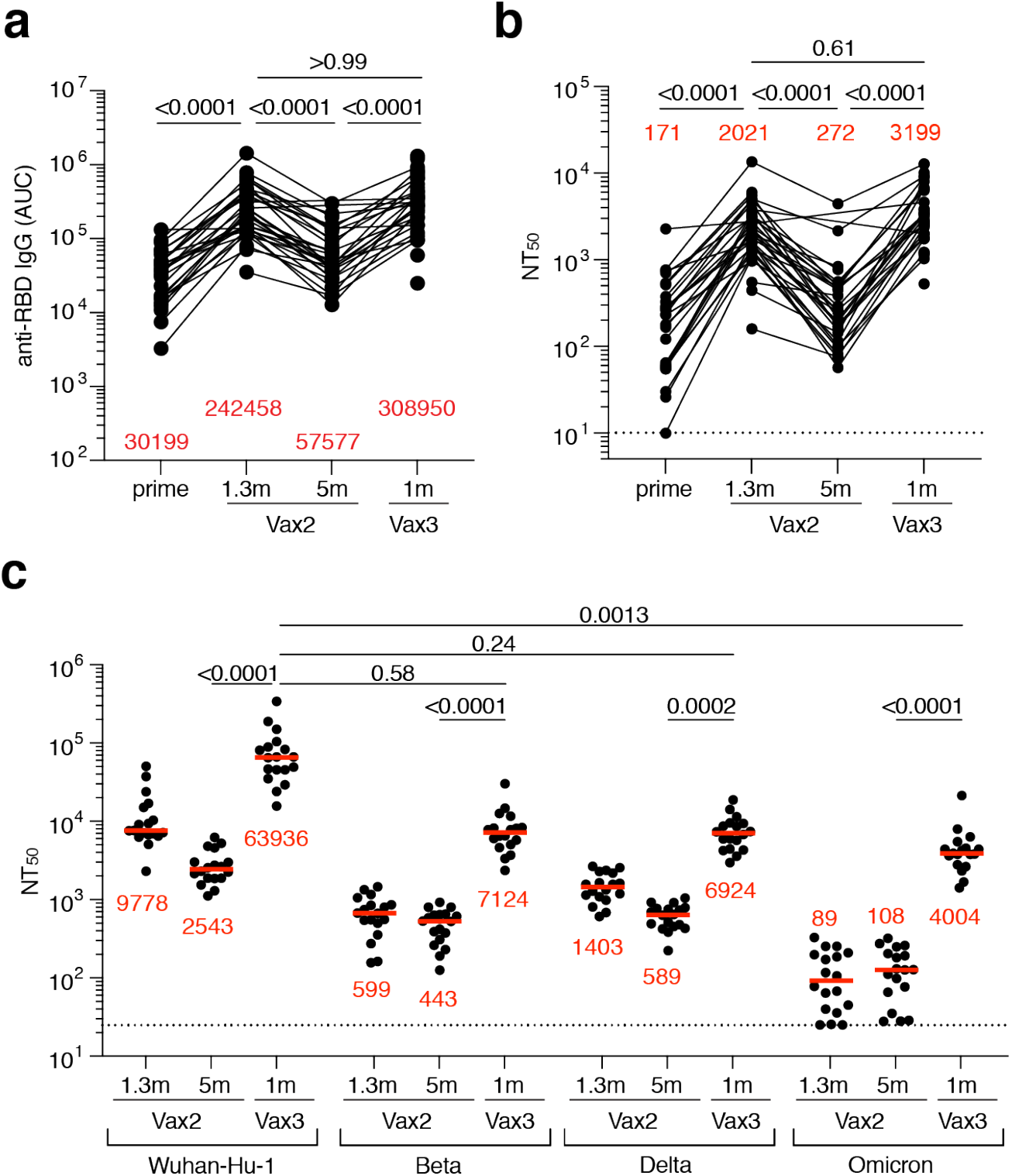
Plasma ELISAs and neutralizing activity. **a**, Graph shows area under the curve (AUC) for plasma IgG antibody binding to SARS-CoV-2 RBD after prime^10^, 1.3 months (m) and 5 months (m) post-second vaccination (Vax2)^9,10^, and 1 month after third vaccination booster (Vax3) for n=43 samples. Lines connect longitudinal samples. **b**, Graph shows anti-SARS-CoV-2 NT50s of plasma measured by a SARS-CoV-2 pseudotype virus neutralization assay using wild-type (Wuhan Hu-1^33^) SARS-CoV-2 pseudovirus^18,34^ in plasma samples shown in panel **a. c**, Plasma neutralizing activity against indicated SARS-CoV-2 variants of interest/concern for n=15 randomly selected samples. Wuhan-Hu-1 and Omicron NT50 values are derived from ^7^. See Methods for a list of all substitutions/deletions/insertions in the spike variants. All experiments were performed at least in duplicate. Red bars and values in **a, b**, and **c** represent geometric mean values. Statistical significance in **a-b** was determined by two-tailed Kruskal-Wallis test with subsequent Dunn’s multiple comparisons. Statistical significance in **c** was determined by Friedman-test with subsequent Dunn’s multiple comparisons.

Plasma neutralizing activity in 43 participants was measured using HIV-1 pseudotyped with the Wuhan-Hu-1 SARS-CoV-2 spike protein^9,10^ (Fig. 1b and Supplementary Table 1). Following a 7.4-fold decrease in neutralizing titers between 1.3- and 5-months after the 2^nd^ vaccine dose, administration of a 3^rd^ vaccine dose boosted neutralizing titers 11.8-fold resulting in a geometric mean half-maximal neutralizing titer (NT_50_) of 3,199 against Wuhan-Hu-1 (Fig. 1b). Plasma neutralizing antibodies elicited by mRNA vaccination are more potent against Wuhan-Hu-1 than variants^9,10^. Consistent with prior reports^3,7,11–13^, the 3^rd^ vaccine dose significantly boosts geometric mean NT_50_s 16-fold, 12-fold and 37-fold for the Beta, Delta and Omicron variant, respectively. The level of activity against the Beta and Delta variants was not significantly different than against Wuhan-Hu-1 while the activity against Omicron was 16-fold lower than against Wuhan-Hu-1 (p=0.58, p=0.24 and p=0.0013, respectively. Fig. 1c). Given the correlation between neutralizing antibody levels and protection from infection^14,15^, the reduced activity against Omicron in 3^rd^ dose vaccine recipients is likely to explain why vaccinees remain particularly susceptible to infection by this variant.

### Memory B cells

Under physiologic conditions memory B cells produce little if any secreted antibody. However, when challenged with antigen as in a breakthrough infection, these cells undergo clonal expansion and produce antibody secreting plasma cells, memory and germinal center B cells^16^. To examine the effects of the 3^rd^ vaccine dose on the memory compartment in our longitudinal cohort we performed flow cytometry experiments using phycoerythrin (PE) and Alexa Fluor 647 (AF647) labeled RBDs (Fig. 2a and Extended data Fig. 2). Individuals that received a 3^rd^ vaccine dose developed significantly increased numbers of RBD-binding memory cells compared to the 2^nd^ dose or naturally infected individuals^9,10,17^ (Fig. 2a and b). The number of memory cells produced after the 3^rd^ dose was also higher than for vaccinated convalescent individuals but did not reach significance (p=0.08, Fig 2b). An increased proportion of memory B cells circulating after the 3^rd^ dose expressed IgG and lower levels of CD71 suggesting that germinal center-derived memory B cells dominate this compartment (Extended data Fig. 2c).

**Fig. 2:**
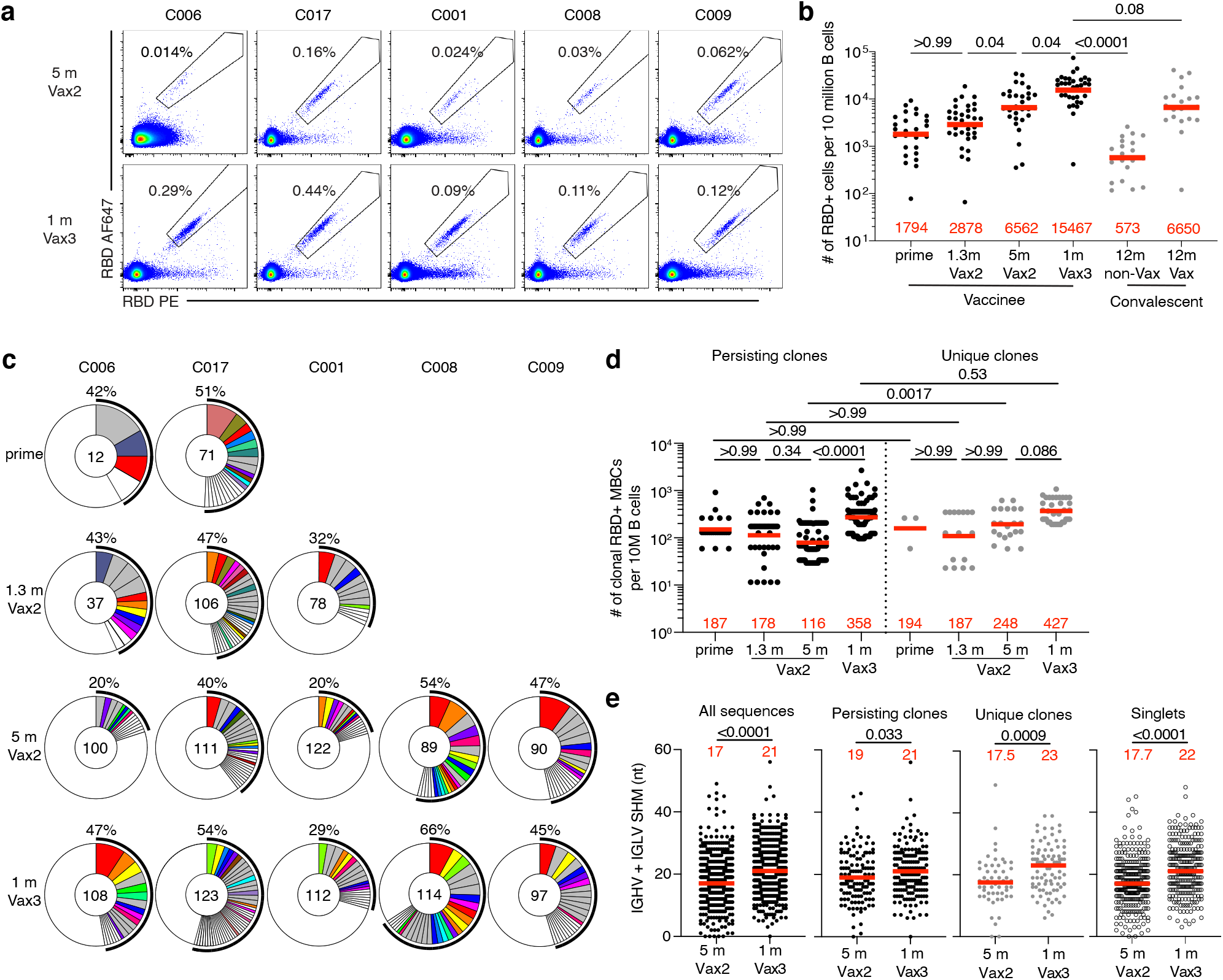
Anti-SARS-CoV-2 RBD memory B cells after third vaccination. **a**, Representative flow cytometry plots showing dual AlexaFluor-647-RBD and PE-RBD-binding, single-cell sorted B cells from 5 individuals 5 months after Vax2^10^ and 1 month after the 3^rd^ vaccine dose (Vax3). Gating strategy is shown in Extended Data Fig. 2. Percentage of RBD-specific B cells is indicated. **b**, Graph summarizing the number of Wuhan-Hu-1 RBD-specific memory B cells (MBCs) per 10 million B cells after prime^9,10^, 1.3- and 5-months after Vax2^9,10^, and 1 month after the 3^rd^ vaccine dose (n=43), compared to the number of RBD-specific MBCs detected in convalescent infected individuals 12-months after infection with or without later vaccination^17^ (shown here in grey). **c**, Pie charts show the distribution of IgG antibody sequences obtained from memory B cells from 5 individuals after prime^10^, 1.3-months and 5-months post-Vax2^9,10^, and 1 month after the 3^rd^ vaccine dose. Time points indicated to the left of the charts. The number inside the circle indicates the number of sequences analyzed for the individual denoted above the circle. Pie slice size is proportional to the number of clonally related sequences. The black outline and associated numbers indicate the percentage of clonal sequences detected at each time point. Colored slices indicate persisting clones (same *IGHV* and *IGLV* genes, with highly similar CDR3s) found at more than one timepoint within the same individual. Grey slices indicate clones unique to the timepoint. White slices indicate repeating sequences isolated only once per time point. **d**, Graph shows the number of clonal RBD-specific MBCs per 10 million B cells. Each dot represents one clone illustrated in Fig. 2c. Left panel (black dots) represent persisting clones. Right panel (grey dots) represent time point unique clones. **e**, Number of nucleotide somatic hypermutations (SHM) in *IGHV* + *IGLV* in all sequences detected 5 months after Vax2^10^ or 1 month after Vax3, compared to SHM in *IGHV* + *IGLV* of sequences from persisting clones, unique clones, and singlets. Red bars and numbers in **b**, and **d**, represent geometric mean value at each time point, and in **e**, represent median values. Statistic difference in **b**, and **d**, was determined by determined by two-tailed Kruskal Wallis test with subsequent Dunn’s multiple comparisons, and in **e**, by two-tailed Mann-Whitney test.

We obtained 1370 paired antibody sequences from 5 individuals who were sampled 5 months after the 2^nd^ and 1 month after the 3^rd^ vaccine dose. Two and 3 out of those participants were additionally sampled 2.5 weeks after the first dose and 1.3 months after the second dose, respectively (^9,10^, Fig. 2c, Supplementary Table 2). After the 3^rd^ vaccine dose all individuals examined showed expanded clones of memory B cells (Fig. 2c). Like earlier time points there was over-representation of VH3-30, VH3-53 and VH4-31 genes (^9,10^ and Extended data Fig. 3). Thus, there is a persistent bias in IGVH gene representation in memory which is common to most individuals.

Expanded clones of memory cells accounted for 33% and 47% of the repertoire 5 months after the 2^nd^ and 1 month after the 3^rd^ dose, respectively (Fig. 2c and Extended data Fig. 4a). The relative increase in clonality was due in part to an average 3.1-fold expansion of persisting anti-RBD-specific memory B cells (p<0.0001, Fig. 2d). Consistent with the relatively modest number of additional cell divisions by persisting clones, they accumulated on average only 2 additional somatic hypermutations making it unlikely that the additional clonal expansion required further germinal center residence^16^ (Fig. 2e and Extended data Fig. 4b).

There was also a more modest 1.7-fold increase in the number of newly emerging unique clones of memory cells after the 3^rd^ dose that did not reach statistical significance (p=0.09) (Fig. 2d). These cells were more mutated than the unique clones present 5 months after the 2^nd^ vaccine dose as were antibodies that were represented only once (singlets). In both cases the numbers of somatic mutations were significantly greater than at 5 months after the 2^nd^ dose indicating persisting evolution and cell division (p=0.0009 and p<0.0001, respectively. Fig. 2e and Extended data Fig. 4). In conclusion, the 3^rd^ mRNA vaccine dose is associated with expansion and further evolution of the memory B cell compartment.

### Monoclonal antibodies

472 monoclonal antibodies obtained from different time points were expressed and tested by ELISA, 459 bound to Wuhan-Hu-1 RBD indicating the high efficiency of the RBD-specific memory B cell isolation method employed here (Extended data Fig. 5 and Supplementary Table 3). 191 antibodies obtained after the 3^rd^ vaccine dose were compared to 34 isolated after the prime; 79 and 168 isolated 1.3 and 5 months after the 2^nd^ vaccine dose (Vax2-1.3m and Vax2-5m), respectively. The geometric mean ELISA half-maximal concentration (EC_50_) of the RBD-binding antibodies was 4.4, 3.8, 2.9 and 3.5 ng/ml for antibodies isolated at the prime, Vax2-1.3-months, Vax2-5-months and Vax3-1-month timepoints, respectively (Extended data Fig. 5a and Supplementary Table 3). Overall, there was no significant change in binding over time or the number of vaccine doses. This was true for all antibodies combined, as well as for persisting clones, unique clones that could only be detected at a single timepoint, and single antibodies (Extended data Fig. 5a-c).

All 459 RBD-binding antibodies were subjected to a SARS-CoV-2 pseudotype neutralization assay based on the Wuhan-Hu-1 SARS-CoV-2 spike^9,10^. Between 1.3- and 5-months after the 2^nd^ vaccine dose antibody potency improved but did not reach statistical significance (IC_50_ 290 vs. 182, p=0.60 Fig 3a). There was additional improvement after the 3^rd^ vaccine dose (IC_50_ 182 vs. 111, p=0.049 Fig. 3a). The overall improvement between equivalent time points after the 2^nd^ and the 3^rd^ dose, from IC_50_ 290 ng/ml to 111 ng/ml was highly significant (p=0.0023, Fig. 3a and Supplementary Table 3). Notably, the potency of antibodies isolated after the 3^rd^ dose, approximately 10 months (293 (223-448) days) after the prime-dose, was indistinguishable from antibodies isolated from convalescent vaccinated individuals 12 months after infection (p=0.69, Fig. 3a) ^**17-19**^. The improved neutralizing activity was most evident among unique clones with a dramatic change in IC_50_ from 323 to 67ng/ml, p=0.034 (Fig. 3b and Supplementary Table 3). Persisting clones also showed improved neutralizing activity after the 3^rd^ dose (p=0.043, Fig. 3b) and a trend to improved neutralizing activity was evident among single antibodies but this did not reach statistical significance (Fig. 3b, Extended data Fig. 5d and Supplementary Table 3 and 4). In all cases, the relative potency of the antibodies isolated 1 month after the 3^rd^ dose was similar to the antibodies isolated from convalescent vaccinated individuals 12 months after infection (Fig. 3a and b). Taken together, there is a significant improvement in the neutralizing potency of the antibodies expressed in the memory B cell compartment 1 month after administration of the 3^rd^ mRNA vaccine dose compared to 1.3 months after the 2^nd^ dose. Newly detected singlets and clones of expanded memory B cells account for most of the improvement in neutralizing activity between 5 months after 2^nd^ dose and 1 month after the 3^rd^ dose.

**Fig. 3:**
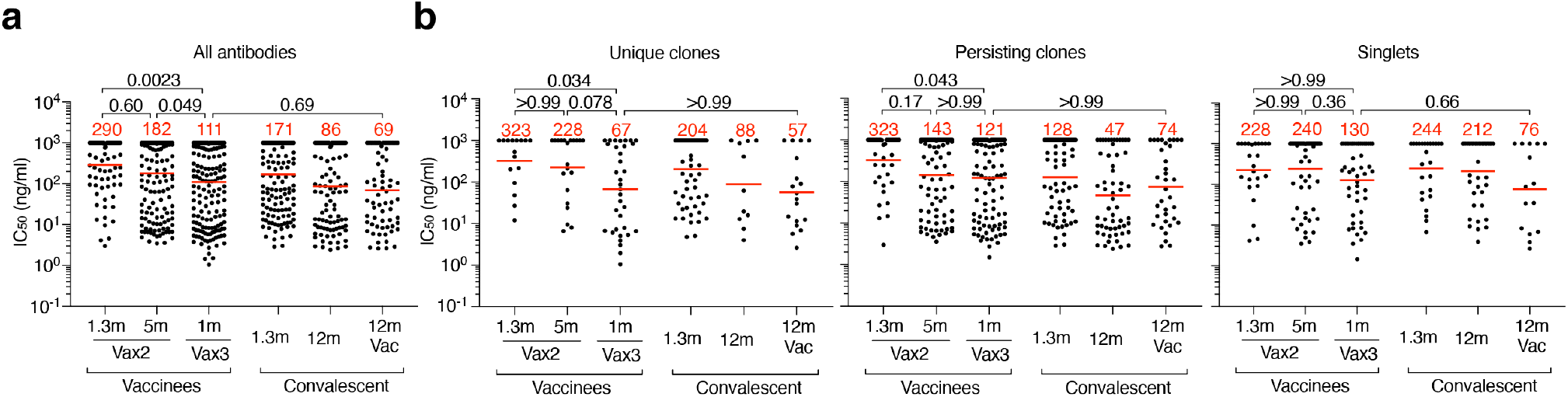
Anti-SARS-CoV-2 RBD monoclonal antibodies. **a-b**, Graphs show anti-SARS-CoV-2 neutralizing activity of monoclonal antibodies measured by a SARS-CoV-2 pseudotype virus neutralization assay using wild-type (Wuhan Hu-1^33^) SARS-CoV-2 pseudovirus^18,34^. IC50 values for all antibodies (**a**), unique clones, persisting clones, and singlets (**b**). Antibodies were from vaccinated individuals 1.3 and 5 months after the 2^nd^ vaccine dose (1.3m-Vax2 and 5m-Vax2, respectively)^9,10^; 1 month after the 3^rd^ vaccination (1m-Vax3): convalescent individuals 1.3 months^18^, or 12 months^17^ after infection or vaccinated convalescent individuals 12 months after infection. Each dot represents one antibody, where 459 total antibodies were tested including the 325 reported herein (Supplementary Table 4), and 134 previously reported^10^. Red bars and numbers indicate geometric mean values. Statistical significance was determined by two-tailed Kruskal Wallis test with subsequent Dunn’s multiple comparisons. All experiments were performed at least twice.

### Epitopes and Neutralization Breadth

The majority of the anti-RBD neutralizing antibodies obtained from vaccinated individuals after the 2^nd^ vaccine dose belong to class 1 and 2 that target a region overlapping with the ACE2 binding site ^20,21^ (Fig. 4a). These antibodies are generally more potent than class 3 and 4 antibodies that target the more conserved base of the RBD and do not directly interfere with ACE2 binding (^17^, Fig. 4a and Extended data Fig. 6). Whereas class 1 and 2 antibodies that develop early are susceptible to mutations in and around the ACE2 binding site found in many of the variants of concern, evolved versions of the same antibodies can be resistant^17,22^. Based on structural information and sequence conservation among betacoronaviruses, antibodies that span class 3 or 4 and either class 1 or 2 could be broadly active (Fig. 4b and Extended data Fig. 6).

**Fig. 4:**
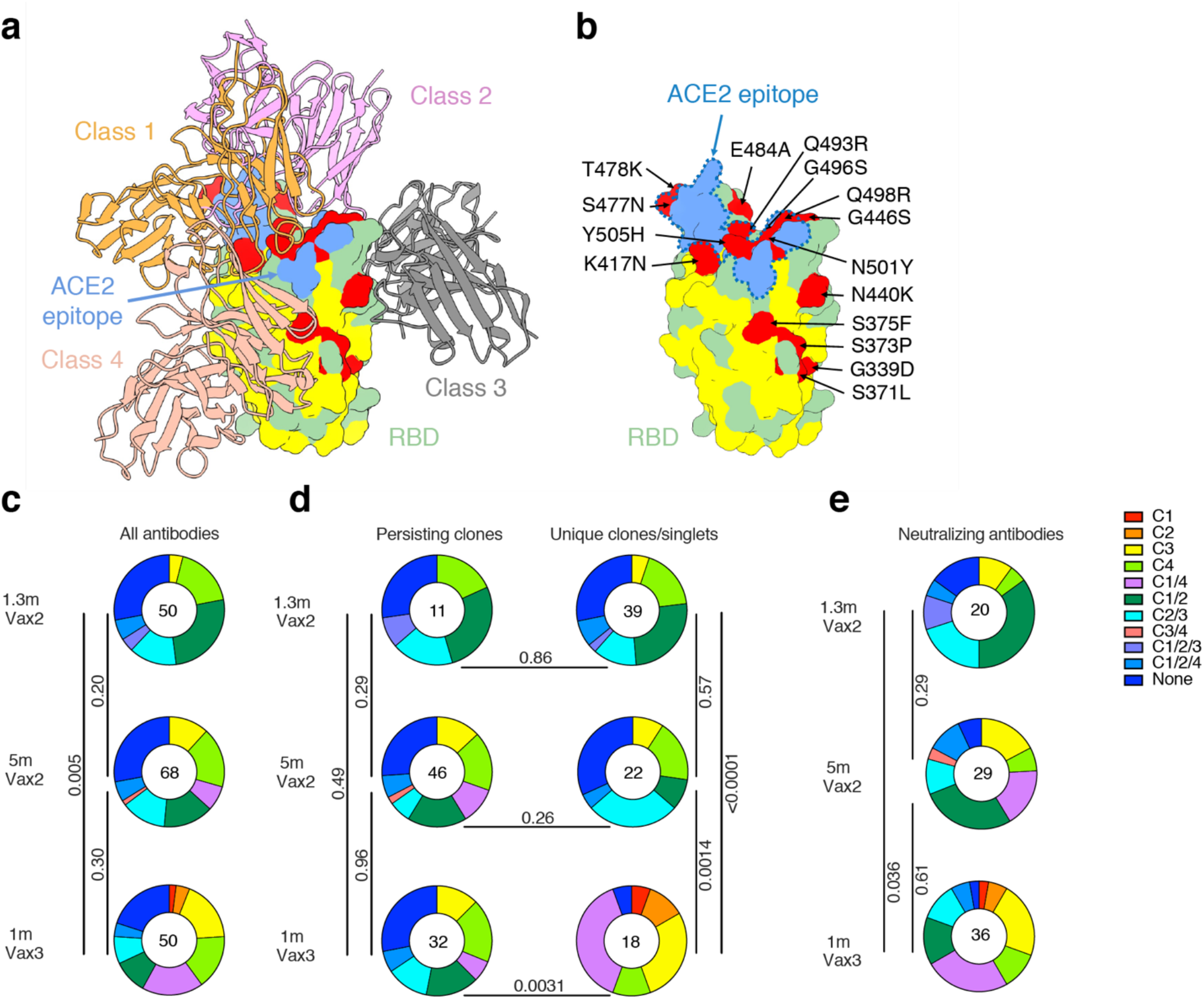
Epitope mapping. **a**, Diagram represents binding poses of antibodies used in BLI competition experiments on the RBD epitope. Class 1 antibody (C105, PDB:6XCM) was shown in orange, class 2 antibody (C144, PDB:7K90) was shown in pink, class 3 antibody (C135 PDB:7K8Z) was shown in gray, and class 4 antibody (C118, PDB:7RKS) was shown in light coral 18,20. ACE2 epitope of Omicron variant was shown in blue. Omicron mutations were shown in red. The most conserved residues calculated by the ConSurf Database were shown in yellow (related to Extended data Fig. 6). **b**, RBD in **a** was enlarged. ACE2 epitope of Omicron variant was indicated by blue dashed lines, and Omicron mutations were labeled. **c-e**, Results of epitope mapping performed by competition BLI. Pie charts show the distribution of the antibody classes among all RBD binding antibodies (**c**), RBD binding antibodies from persisting clones or unique clones or singlets (**d**), or neutralizing antibodies against Wuhan-Hu-1 (**e**) obtained 1.3- and 5-months after Vax2^9,10^, and 1 month after 3^rd^ vaccine dose (Vax3). Statistical significance was determined using a two-tailed Chi-square test.

To examine epitopes targeted by RBD-binding antibodies after the 3^rd^ vaccine dose, we performed BLI experiments in which a preformed antibody-RBD complex was exposed to a second antibody targeting one of four classes of structurally defined epitopes (C105 as Class 1; C144 as Class 2, C135 as Class 3 and C118 as Class 1/4 ^18,20^) (Fig. 4a). 168 random RBD binding antibodies were tested among which 20, 29, and 36 neutralized with IC_50_s lower than 1000 ng/ml from 1.3 and 5-months after the 2^nd^ and 1 month after the 3^rd^ vaccine dose respectively. As might be expected the largest group of RBD binding antibodies obtained after the 2^nd^ vaccine dose belonged to class 1/2 (Fig. 4c). Although the overall distribution of antibody classes that bind to RBD did not change significantly between 1.3 and 5-months after the 2^nd^ dose, the relative representation of class 1 and 2 antibodies decreased (Fig. 4c). This trend continued after the 3^rd^ vaccine dose with increased representation of RBD binding antibodies in class 1/4 and 3 resulting in a significant difference in the epitope distribution among RBD-binding antibodies between the early time points after the 2^nd^ and the 3^rd^ dose (p=0.005, Fig. 4c). As expected, these differences can be accounted for primarily by the emergence of new clones and singlets after the 3^rd^ vaccine dose (Fig. 4d). Similar results were found when considering the neutralizing antibodies with initial dominance of class 1/2 and increasing representation of class 1/4 and 3 over time (Fig. 4e).

The neutralizing breadth of antibodies elicited by infection increased significantly after 5 months ^17,19,22^. There was also a trend to increased breadth that did not reach statistical significance 5 months after the 2^nd^ dose of an mRNA vaccine^10^. To determine whether neutralizing antibodies in clones that persisted from 5 months after the 2^nd^ to 1 month after the 3^rd^ dose develop increased breadth, we compared 18 antibody pairs. Neutralizing activity was measured against a panel of SARS-CoV-2 pseudoviruses harboring RBD amino acid substitutions representative of SARS-CoV-2 variants including Delta and Omicron (Fig. 5a). The clonal pairs were dominated by antibodies belonging to class 1/2, 2/3 and 3, as determined by BLI (Fig 5a). 15 out of 18 antibody pairs neutralized the pseudovirus carrying the Delta RBD-amino acid substitutions at low antibody concentrations at both time points, with IC_50_ values ranging from 1-154 ng/ml (Fig. 5a). While the Omicron pseudovirus showed the highest degree of neutralization resistance, 11 out of 18 antibodies isolated 1 month after the 3^rd^ dose neutralized this virus, 9 of those at IC_50_s below 120 ng/ml (Fig. 5a). Most antibody pairs isolated before and after the 3^rd^ vaccine dose showed exceptionally broad neutralization and there was little change in antibody breadth within the analyzed pairs (Fig. 5a).

**Fig. 5:**
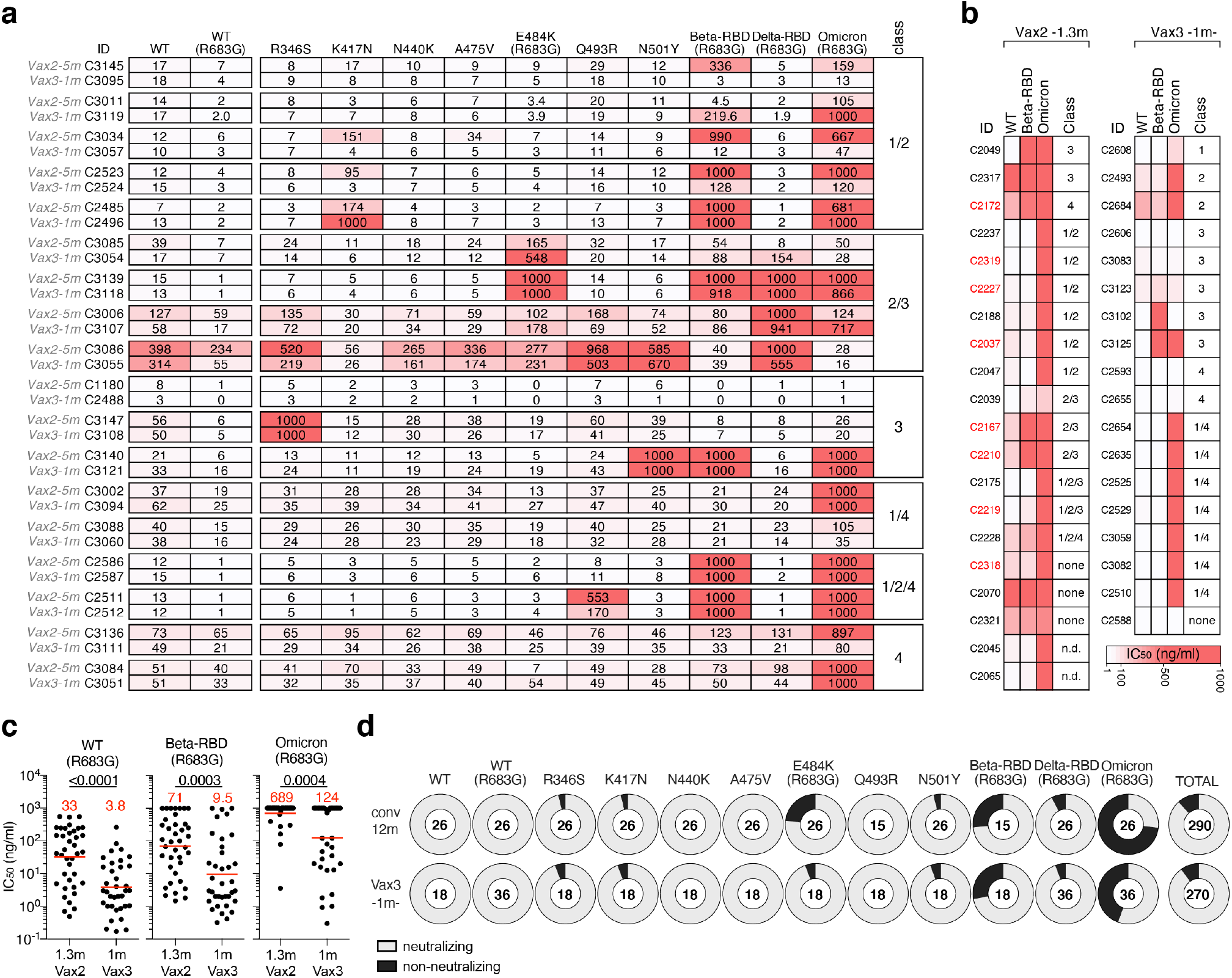
Breadth. **a-b** Heat-maps show IC50s of clonal pairs of antibodies detected 5 months after the 2^nd^ vaccination (Vax2-5m) persisting 1 month after the 3^rd^ dose (Vax3-1m) (**a**) and clones and singlets found 1.3 months after the 2^nd^ (Vax2-1.3m) and uniquely 1 month after the 3^rd^ (Vax3-1m) vaccine dose (**b**), against indicated mutant and variant SARS-CoV-2 pseudoviruses listed across the top. Beta-RBD and Delta-RBD indicate the K417N/E484K/N501Y and L452R/T478K SARS-CoV-2 spikes, respectively. Heatmap ranging from 0.1-1000 ng/ml in white to red. Antibody classes in **a** and **b** were determined by competition BLI. **c**, graphs show neutralization activity of antibodies shown in **a** and **b** against WT, Beta-RBD (L452R/T478K) and Omicron, comparing 1.3-month Vax2 and 1-month Vax3 timepoints. Red bars and numbers indicate geometric mean values. Statistical significance was determined using two-tailed Mann-Whitney test. **d**, Ring plots show fraction of neutralizing (IC50<1000ng/ml) and non-neutralizing (IC50>1000 ng/ml) antibodies in light and dark grey, respectively, for indicated SARS-CoV-2 pseudoviruses. Number in inner circles indicates number of antibodies tested.

We extended the analysis to compare the activity of antibodies present in memory cells found 1.3 months after the 2^nd^ and unique to 1 month after the 3^rd^ vaccine dose. The antibodies were tested against viruses pseudotyped with spike proteins containing the RBD of Wuhan-Hu-1, Delta and Omicron (Fig. 5b). We found that the proportion of Omicron-neutralizing antibodies increased from 15% after the 2^nd^ dose to 50% among the unique antibodies found after the 3^rd^ dose (p=0.035, Fisher’s exact test. Fig 5b). Among all antibodies evaluated, the increase in breadth between the 2^nd^ and 3^rd^ vaccine dose was reflected by an increase in potency from 689 to 124 ng/ml IC_50_ against Omicron (p=0.0004, Fig 5c). Similar results were seen for Delta neutralization (Fig. 5c). Thus, memory B cell clones emerging after the 3^rd^ vaccine dose show increasing breadth and potency against pseudoviruses representing variants that were not present in the vaccine.

Finally, we compared the neutralization breadth of 3^rd^ dose vaccine-elicited antibodies, as measured approximately 10 months (292 (223-448) days) after the prime dose, with antibodies we obtained from a cohort of convalescent unvaccinated individuals 12 months after infection (^17–19^ and Fig. 5d). The two groups of antibodies are equally and remarkably broad. 92% and 94% of the convalescent and 3^rd^ dose antibodies neutralized pseudoviruses carrying the Beta-RBD and 27% and 56%, respectively, neutralized Omicron. Thus, 3^rd^ dose vaccine-elicited antibodies are at least as broad as those elicited by infection (Fig. 5d).

## Discussion

Memory B cells can develop from the germinal center or directly from a germinal center independent activated B cell compartment^16^. B cells residing in germinal centers undergo multiple rounds of division, mutation and selection, whereas those in the activated compartment undergo only a limited number of divisions and carry fewer mutations^16^. Both pathways remain active throughout the immune response^23,24^. Our data indicate that the 3^rd^ dose of mRNA vaccines against SARS-CoV-2 expands persisting clones of memory B cells through the germinal center independent compartment because these cells show limited clonal expansion and accumulate a small number of additional mutations. In addition, however, the 3^rd^ dose elicits a cohort of previously undetected clones that carry mutations indicative of germinal center residence. The later differ from the persistent clones in that they appear to target more conserved regions of the RBD. Several different mechanisms could account for the antigenic shift including epitope masking by the high affinity antibodies elicited by earlier vaccine doses that primarily target the less conserved receptor binding domain of the RBD^20,21,25^.

Passively administered antibodies are protective against SARS-CoV-2 infection and can also prevent serious disease if provided early^26–30^. The 3^rd^ dose of mRNA vaccines boosts plasma antibody responses to multiple SARS-CoV-2 variants including Omicron, but the levels are insufficient to prevent breakthrough infection in many individuals^2,3^. The 3^rd^ dose also elicits increased number of memory B cells that express more potent and broader antibodies. These cells do not appear to contribute to circulating plasma antibody levels, but upon challenge with antigen in the form of a vaccine or infection, they produce large amounts of antibodies within 3-5 days^31^. Passive administration of antibodies within this same time window prevents the most serious consequences of infection^26,29,32^. Thus, rapid recall by a diversified and expanded memory B cell compartment is likely to be one of the key mechanisms that contribute to the enhanced protection against severe disease by a 3^rd^ mRNA vaccine dose.

## Methods

### Study participants

Participants were healthy volunteers who had previously received the initial two-dose regimen of either the Moderna (mRNA-1273) or Pfizer-BioNTech (BNT162b2) mRNA vaccines against the wildtype (Wuhan-Hu-1) strain of the severe acute respiratory syndrome coronavirus 2 (SARS-CoV-2). For this study, participants were recruited for serial blood donations at the Rockefeller University Hospital in New York between January 21, 2021, and December 14, 2021. The majority of participants (n=32) were follow-ups from a longitudinal cohort that we previously reported on^9,10^, while a smaller subgroup of individuals (n=11) was de novo recruited for this study (for details see Supplementary Table 1). Eligible participants (n=43) were healthy adults with no history of infection with SARS-CoV-2 during or prior to the observation period (as determined by clinical history and confirmed through serology testing) who had received two doses of one of the two currently approved SARS-CoV-2 mRNA vaccines, Moderna (mRNA-1273) or Pfizer-BioNTech (BNT162b2), and this also included a subgroup of individuals (n=34) who had received a third vaccine dose. The specifics of each participant’s vaccination regimen were at the discretion of the individual and their health care provider consistent with current dosing and interval guidelines and, as such, not influenced by their participation in our study. Exclusion criteria included incomplete vaccination status (defined as less than 2 doses), presence of clinical signs and symptoms suggestive of acute infection with or a positive reverse transcription polymerase chain reaction (RT-PCR) results for SARS-CoV-2 in saliva, or a positive (coronavirus disease 2019) COVID-19 serology. Participants presented to the Rockefeller University Hospital for blood sample collection and were asked to provide details of their vaccination regimen, possible side effects, comorbidities and possible COVID-19 history. Clinical data collection and management were carried out using the software iRIS by iMedRIS (v. 11.02). All participants provided written informed consent before participation in the study and the study was conducted in accordance with Good Clinical Practice. The study was performed in compliance with all relevant ethical regulations and the protocol (DRO-1006) for studies with human participants was approved by the Institutional Review Board of the Rockefeller University. For detailed participant characteristics see Supplementary Table 1.

### Blood samples processing and storage

Peripheral Blood Mononuclear Cells (PBMCs) obtained from samples collected at Rockefeller University were purified as previously reported by gradient centrifugation and stored in liquid nitrogen in the presence of Fetal Calf Serum (FCS) and Dimethylsulfoxide (DMSO)^18,19^. Heparinized plasma and serum samples were aliquoted and stored at −20°C or less. Prior to experiments, aliquots of plasma samples were heat-inactivated (56°C for 1 hour) and then stored at 4°C.

### ELISAs

Enzyme-Linked Immunosorbent Assays (ELISAs)^35,36^ to evaluate antibodies binding to SARS-CoV-2 RBD were performed by coating of high-binding 96-half-well plates (Corning 3690) with 50 μl per well of a 1μg/ml protein solution in Phosphate-buffered Saline (PBS) overnight at 4°C. Plates were washed 6 times with washing buffer (1× PBS with 0.05% Tween-20 (Sigma-Aldrich)) and incubated with 170 μl per well blocking buffer (1× PBS with 2% BSA and 0.05% Tween-20 (Sigma)) for 1 hour at room temperature. Immediately after blocking, monoclonal antibodies or plasma samples were added in PBS and incubated for 1 hour at room temperature. Plasma samples were assayed at a 1:66 starting dilution and 10 additional threefold serial dilutions. Monoclonal antibodies were tested at 10 μg/ml starting concentration and 10 additional fourfold serial dilutions. Plates were washed 6 times with washing buffer and then incubated with anti-human IgG, IgM or IgA secondary antibody conjugated to horseradish peroxidase (HRP) (Jackson Immuno Research 109-036-088 109-035-129 and Sigma A0295) in blocking buffer at a 1:5,000 dilution (IgM and IgG) or 1:3,000 dilution (IgA). Plates were developed by addition of the HRP substrate, 3,3’,5,5’- Tetramethylbenzidine (TMB) (ThermoFisher) for 10 minutes (plasma samples) or 4 minutes (monoclonal antibodies). The developing reaction was stopped by adding 50 μl of 1 M H_2_SO_4_ and absorbance was measured at 450 nm with an ELISA microplate reader (FluoStar Omega, BMG Labtech) with Omega and Omega MARS software for analysis. For plasma samples, a positive control (plasma from participant COV72, diluted 66.6-fold and ten additional threefold serial dilutions in PBS) was added to every assay plate for normalization. The average of its signal was used for normalization of all the other values on the same plate with Excel software before calculating the area under the curve using Prism V9.1(GraphPad). Negative controls of pre-pandemic plasma samples from healthy donors were used for validation (for more details please see^18^). For monoclonal antibodies, the ELISA half-maximal concentration (EC50) was determined using four-parameter nonlinear regression (GraphPad Prism V9.1). EC50s above 1000 ng/mL were considered non-binders.

### Proteins

The mammalian expression vector encoding the Receptor Binding-Domain (RBD) of SARS-CoV-2 (GenBank MN985325.1; Spike (S) protein residues 319-539) was previously described^37^.

### SARS-CoV-2 pseudotyped reporter virus

A panel of plasmids expressing RBD-mutant SARS-CoV-2 spike proteins in the context of pSARS-CoV-2-S _Δ19_ has been described^9,10,22,38^. Variant pseudoviruses resembling SARS-CoV-2 variants Beta (B.1.351), B.1.526, Delta (B.1.617.2) and Omicron (B.1.1.529) habe been descrived before ^7,10,17^ and were generated by introduction of substitutions using synthetic gene fragments (IDT) or overlap extension PCR mediated mutagenesis and Gibson assembly. Specifically, the variant-specific deletions and substitutions introduced were:

Beta: D80A, D215G, L242H, R246I, K417N, E484K, N501Y, D614G, A701V

DeltaB.1.617.2: T19R, Δ156-158, L452R, T478K, D614G, P681R, D950N

Omicron: A67V, Δ69-70, T95I, G142D, Δ143-145, Δ211, L212I, ins214EPE, G339D, S371L, S373P, S375F, K417N, N440K, G446S, S477N, T478K, E484A, Q493K, G496S, Q498R, N501Y, Y505H, T547K, D614G, H655Y, H679K, P681H, N764K, D796Y, N856K, Q954H, N969H, N969K, L981F

The E484K, K417N/E484K/N501Y and L452R/T478K substitution, as well as the deletions/substitutions corresponding to variants of concern listed above were incorporated into a spike protein that also includes the R683G substitution, which disrupts the furin cleaveage site and increases particle infectivity. Neutralizing activity against mutant pseudoviruses were compared to a wildtype (WT) SARS-CoV-2 spike sequence (NC_045512), carrying R683G where appropriate.

SARS-CoV-2 pseudotyped particles were generated as previously described^18,34^. Briefly, 293T (CRL-11268) cells were obtained from ATCC, and the cells were transfected with pNL4-3ΔEnv-nanoluc and pSARS-CoV-2-S_Δ19_, particles were harvested 48 hours post-transfection, filtered and stored at −80°C.

### Pseudotyped virus neutralization assay

Fourfold serially diluted pre-pandemic negative control plasma from healthy donors, plasma from individuals who received mRNA vaccines or monoclonal antibodies were incubated with SARS-CoV-2 pseudotyped virus for 1 hour at 37 °C. The mixture was subsequently incubated with 293T_Ace2_ cells^18^ (for all WT neutralization assays) or HT1080Ace2 cl14 (for all mutant panels and variant neutralization assays) cells^9^ for 48 hours after which cells were washed with PBS and lysed with Luciferase Cell Culture Lysis 5× reagent (Promega). Nanoluc Luciferase activity in lysates was measured using the Nano-Glo Luciferase Assay System (Promega) with the Glomax Navigator (Promega). The relative luminescence units were normalized to those derived from cells infected with SARS-CoV-2 pseudotyped virus in the absence of plasma or monoclonal antibodies. The half-maximal neutralization titers for plasma (NT_50_) or half-maximal and 90% inhibitory concentrations for monoclonal antibodies (IC_50_ and IC_90_) were determined using four-parameter nonlinear regression (least squares regression method without weighting; constraints: top=1, bottom=0) (GraphPad Prism).

### Biotinylation of viral protein for use in flow cytometry

Purified and Avi-tagged SARS-CoV-2 Wuhan-Hu-1 RBD was biotinylated using the Biotin-Protein Ligase-BIRA kit according to manufacturer’s instructions (Avidity) as described before^18^. Ovalbumin (Sigma, A5503-1G) was biotinylated using the EZ-Link Sulfo-NHS-LC-Biotinylation kit according to the manufacturer’s instructions (Thermo Scientific). Biotinylated ovalbumin was conjugated to streptavidin-BV711 for single-cell sorts (BD biosciences, 563262) or to streptavidin-BB515 for phenotyping panel (BD, 564453). RBD was conjugated to streptavidin-PE (BD Biosciences, 554061) and streptavidin-AF647 (Biolegend, 405237)^18^.

### Flow cytometry and single cell sorting

Single-cell sorting by flow cytometry was described previously^18^. Briefly, peripheral blood mononuclear cells were enriched for B cells by negative selection using a pan-B-cell isolation kit according to the manufacturer’s instructions (Miltenyi Biotec, 130-101-638). The enriched B cells were incubated in Flourescence-Activated Cell-sorting (FACS) buffer (1× PBS, 2% FCS, 1 mM ethylenediaminetetraacetic acid (EDTA)) with the following anti-human antibodies (all at 1:200 dilution): anti-CD20-PECy7 (BD Biosciences, 335793), anti-CD3-APC-eFluro 780 (Invitrogen, 47-0037-41), anti-CD8-APC-eFluor 780 (Invitrogen, 47-0086-42), anti-CD16-APC-eFluor 780 (Invitrogen, 47-0168-41), anti-CD14-APC-eFluor 780 (Invitrogen, 47-0149-42), as well as Zombie NIR (BioLegend, 423105) and fluorophore-labeled RBD and ovalbumin (Ova) for 30 min on ice. Single CD3-CD8-CD14-CD16−CD20+Ova−RBD-PE+RBD-AF647+ B cells were sorted into individual wells of 96-well plates containing 4 μl of lysis buffer (0.5× PBS, 10 mM Dithiothreitol (DTT), 3,000 units/ml RNasin Ribonuclease Inhibitors (Promega, N2615) per well using a FACS Aria III and FACSDiva software (Becton Dickinson) for acquisition and FlowJo for analysis. The sorted cells were frozen on dry ice, and then stored at −80 °C or immediately used for subsequent RNA reverse transcription. For B cell phenotype analysis, in addition to above antibodies, B cells were also stained with following anti-human antibodies (all at 1:200 dilution): anti-IgD-BV650 (BD, 740594), anti-CD27-BV786 (BD biosciences, 563327), anti-CD19-BV605 (Biolegend, 302244), anti-CD71-PerCP-Cy5.5 (Biolegend, 334114), anti-IgG-PECF594 (BD, 562538), anti-IgM-AF700 (Biolegend, 314538), anti-IgA-Viogreen (Miltenyi Biotec, 130-113-481).

### Antibody sequencing, cloning and expression

Antibodies were identified and sequenced as described previously^18,39^. In brief, RNA from single cells was reverse-transcribed (SuperScript III Reverse Transcriptase, Invitrogen, 18080-044) and the cDNA was stored at −20 °C or used for subsequent amplification of the variable IGH, IGL and IGK genes by nested PCR and Sanger sequencing. Sequence analysis was performed using MacVector. Amplicons from the first PCR reaction were used as templates for sequence-and ligation-independent cloning into antibody expression vectors. Recombinant monoclonal antibodies were produced and purified as previously described^18^.

### Biolayer interferometry

Biolayer interferometry assays were performed as previously described^18^. Briefly, we used the Octet Red instrument (ForteBio) at 30 °C with shaking at 1,000 r.p.m. Epitope binding assays were performed with protein A biosensor (ForteBio 18-5010), following the manufacturer’s protocol “classical sandwich assay” as follows: (1) Sensor check: sensors immersed 30 sec in buffer alone (buffer ForteBio 18-1105), (2) Capture 1st Ab: sensors immersed 10 min with Ab1 at 10 µg/mL, (3) Baseline: sensors immersed 30 sec in buffer alone, (4) Blocking: sensors immersed 5 min with IgG isotype control at 10 µg/mL. (5) Baseline: sensors immersed 30 sec in buffer alone, (6) Antigen association: sensors immersed 5 min with RBD at 10 µg/mL. (7) Baseline: sensors immersed 30 sec in buffer alone. (8) Association Ab2: sensors immersed 5 min with Ab2 at 10 µg/mL. Curve fitting was performed using the Fortebio Octet Data analysis software (ForteBio).

### Computational analyses of antibody sequences

Antibody sequences were trimmed based on quality and annotated using Igblastn v.1.14. with IMGT domain delineation system. Annotation was performed systematically using Change-O toolkit v.0.4.540^40^. Heavy and light chains derived from the same cell were paired, and clonotypes were assigned based on their V and J genes using in-house R and Perl scripts. All scripts and the data used to process antibody sequences are publicly available on GitHub (https://github.com/stratust/igpipeline/tree/igpipeline2_timepoint_v2).

The frequency distributions of human V genes in anti-SARS-CoV-2 antibodies from this study was compared to 131,284,220 IgH and IgL sequences generated by^41^ and downloaded from cAb-Rep^42^, a database of human shared BCR clonotypes available at https://cab-rep.c2b2.columbia.edu/. Based on the 150 distinct V genes that make up the 1650 analyzed sequences from Ig repertoire of the 5 participants present in this study, we selected the IgH and IgL sequences from the database that are partially coded by the same V genes and counted them according to the constant region. The frequencies shown in Extended Data Fig. 3 are relative to the source and isotype analyzed. We used the two-sided binomial test to check whether the number of sequences belonging to a specific IGHV or IGLV gene in the repertoire is different according to the frequency of the same IgV gene in the database. Adjusted p-values were calculated using the false discovery rate (FDR) correction. Significant differences are denoted with stars.

Nucleotide somatic hypermutation and Complementarity-Determining Region (CDR3) length were determined using in-house R and Perl scripts. For somatic hypermutations, *IGHV* and *IGLV* nucleotide sequences were aligned against their closest germlines using Igblastn and the number of differences were considered nucleotide mutations. The average number of mutations for V genes was calculated by dividing the sum of all nucleotide mutations across all participants by the number of sequences used for the analysis.

## Data presentation

Figures arranged in Adobe Illustrator 2022.

## Data availability statement

Data are provided in Supplementary Tables 1-4. The raw sequencing data and computer scripts associated with Figure 2 have been deposited at Github (https://github.com/stratust/igpipeline/tree/igpipeline2_timepoint_v2). This study also uses data from “A Public Database of Memory and Naive B-Cell Receptor Sequences” (https://doi.org/10.5061/dryad.35ks2), PDB (6VYB and 6NB6), cAb-Rep (https://cab-rep.c2b2.columbia.edu/), Sequence Read Archive (accession SRP010970), and from “High frequency of shared clonotypes in human B cell receptor repertoires” (https://doi.org/10.1038/s41586-019-0934-8).

## Code availability statement

Computer code to process the antibody sequences is available at GitHub (https://github.com/stratust/igpipeline/tree/igpipeline2_timepoint_v2).

## Acknowledgements

We thank all study participants who devoted time to our research, The Rockefeller University Hospital nursing staff and Clinical Research Support Office. We thank all members of the M.C.N. laboratory for helpful discussions, Maša Jankovic for laboratory support and Kristie Gordon for technical assistance with cell-sorting experiments. This work was supported by NIH grant P01-AI138398-S1 (M.C.N.) and 2U19AI111825 (M.C.N.). R37-AI64003 to P.D.B.; R01AI78788 to T.H.. F.M. was supported by the Bulgari Women and Science Fellowship for COVID-19 Research. C.G. was supported by the Robert S. Wennett Post-Doctoral Fellowship, in part by the National Center for Advancing Translational Sciences (National Institutes of Health Clinical and Translational Science Award program, grant UL1 TR001866), and by the Shapiro-Silverberg Fund for the Advancement of Translational Research. P.D.B. and M.C.N. are Howard Hughes Medical Institute Investigators. This article is subject to HHMI’s Open Access to Publications policy. HHMI lab heads have previously granted a nonexclusive CC BY 4.0 license to the public and a sublicensable license to HHMI in their research articles. Pursuant to those licenses, the author-accepted manuscript of this article can be made freely available under a CC BY 4.0 license immediately upon publication.

## Author information

F.M. Z.W. and A.C. contributed equally to this work.

## Author Contributions

F.M., Z.W., A.C., T.H., P.D.B., and M.C.N. conceived, designed, and analyzed the experiments. M. Caskey and C.G. designed clinical protocols. F.M. Z.W., A.C., T.B.T, J.D., E.B., S.Z., R.R., D.S.-B., K.Y., and F.S. carried out experiments. B.J. and A.G., produced antibodies. M.T., K.G.M., I.S., M.D., C.G. and M.C. recruited participants, executed clinical protocols, and processed samples. T.Y.O. and V.R. performed bioinformatic analysis. F.M., Z.W., A.C., T.H., P.D.B., and M.C.N. wrote the manuscript with input from all co-authors.

## Corresponding authors

Correspondence should be addressed to Theodora Hatziioannou, Paul D. Bieniasz, or Michel C. Nussenzweig.

## Extended data Figures

**Extended Data Fig. 1:**
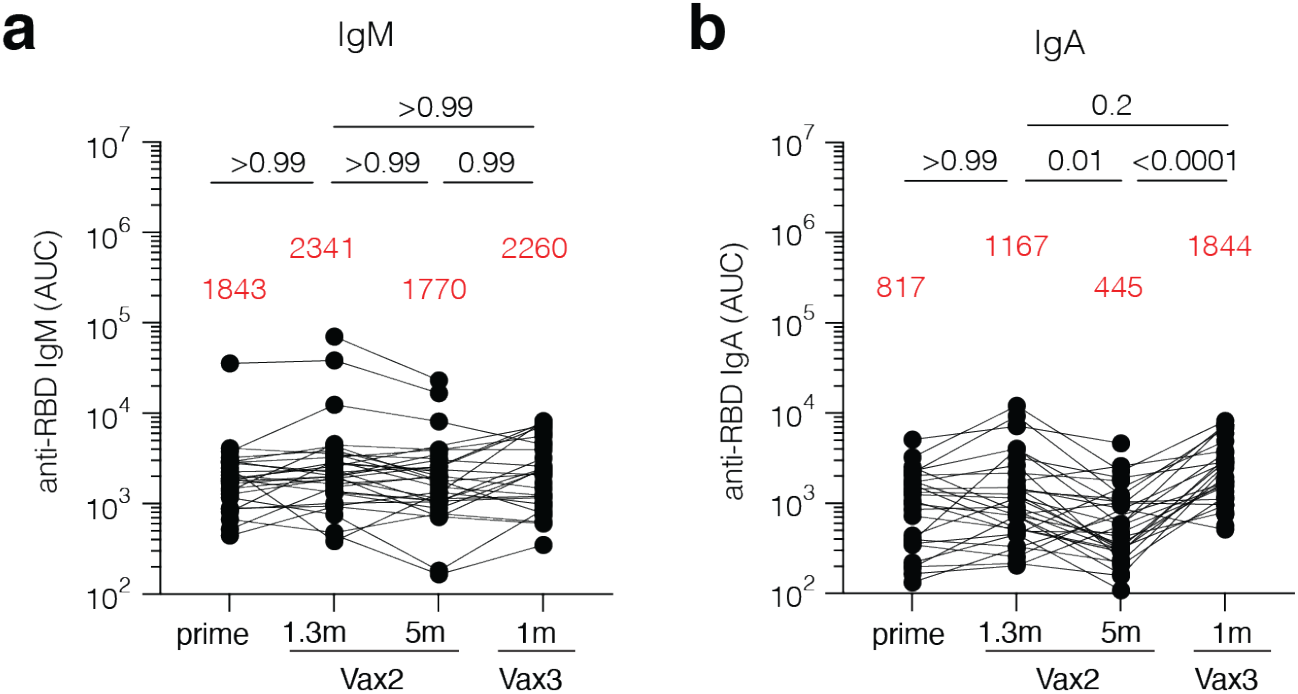
Plasma ELISA. Graph shows area under the curve (AUC) for plasma **a**, IgM and **b**, IgA antibody binding to SARS-CoV-2 RBD after prime^10^, 1.3 months (m) and 5 months (m) after the 2^nd^ vaccine dose (Vax2)^9,10^, and 1 month after the 3^rd^ (Vax3) for n=43 samples. Lines connect longitudinal samples. Red bars and value represent geometric mean values. Statistical significance was determined by two-tailed Kruskal-Wallis test with subsequent Dunn’s multiple comparisons.

**Extended Data Fig. 2:**
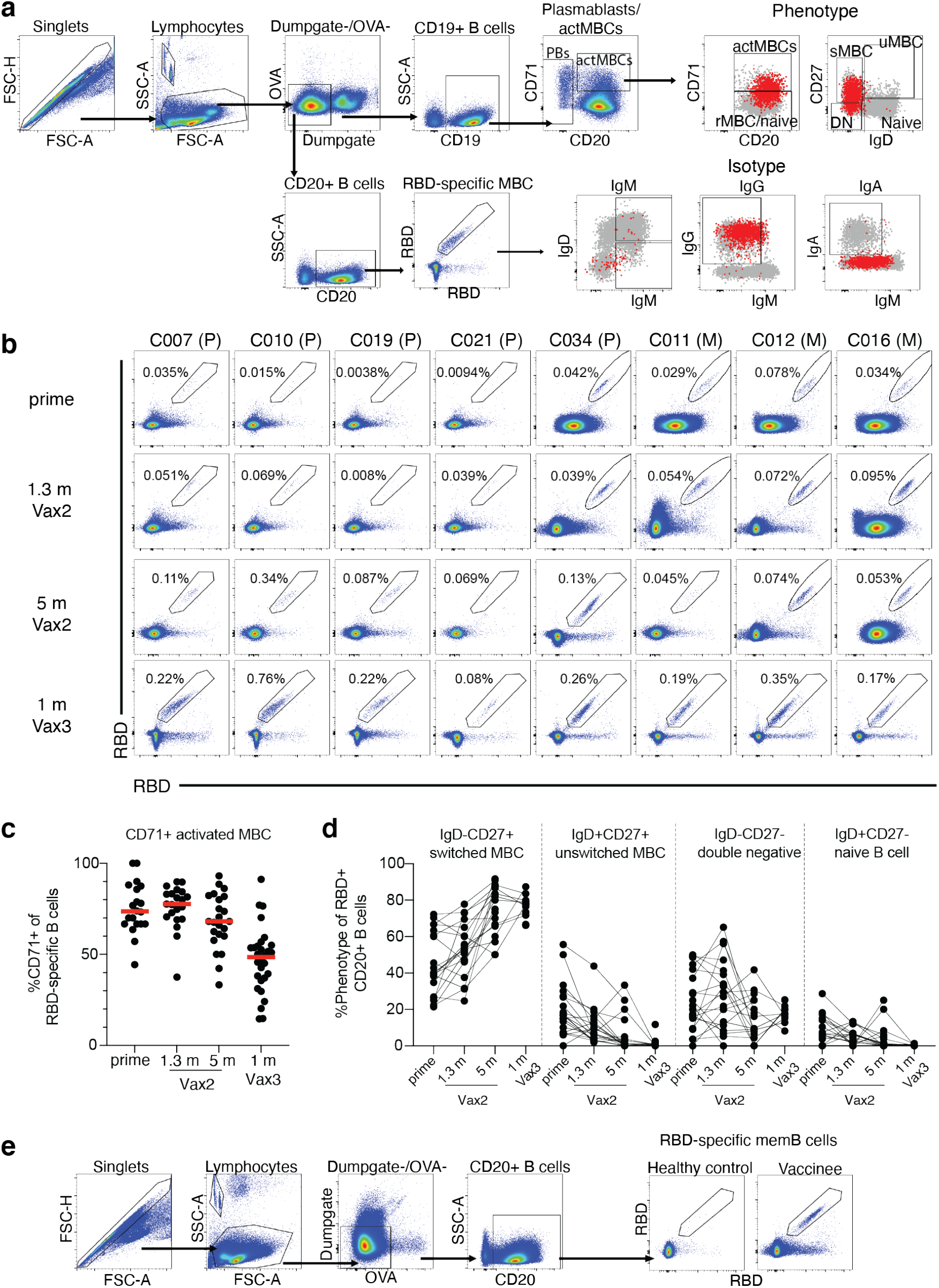
Flow Cytometry. **a**, Gating strategy for phenotyping. Gating was on lymphocytes singlets that were CD19^+^ or CD20^+^ and CD3-CD8-CD16-Ova-. Anti-IgG, IgM, IgA, IgD, CD71 and CD27 antibodies were used for B cell phenotype analysis. Antigen-specific cells were detected based on binding to Wuhan-Hu-1 RBD-PE^+^ and RBD-AF647^+^. **b**, Representative flow cytometry plots of RBD-binding memory B cells in 8 individuals after prime^10^, 1.3- and 5-months post-Vax2^9,10^, and 1 month after Vax3. Time point of sample collection indicated to the left. Pfizer vaccinees indicated by (P) and Moderna by (M) across the top. **c**, Graph showing frequency of RBD-specific MBCs expressing activation marker CD71 over time after vaccination for n=36 samples. Red bar indicated median value. **d**, Graph showing the phenotype of RBD-specific B cells over time, determined to be either switched MBCs (IgD-CD27+), unswitched MBCs (IgD+CD27+), double negative MBCs (IgD-CD27-) or naïve B cells (IgD+CD27+), for n=18 samples. Lines connect longitudinal samples. **f**, Gating strategy for single-cell sorting for CD20+ memory B cells for RBD-PE and RBD-AF647.

**Extended Data Fig. 3:**
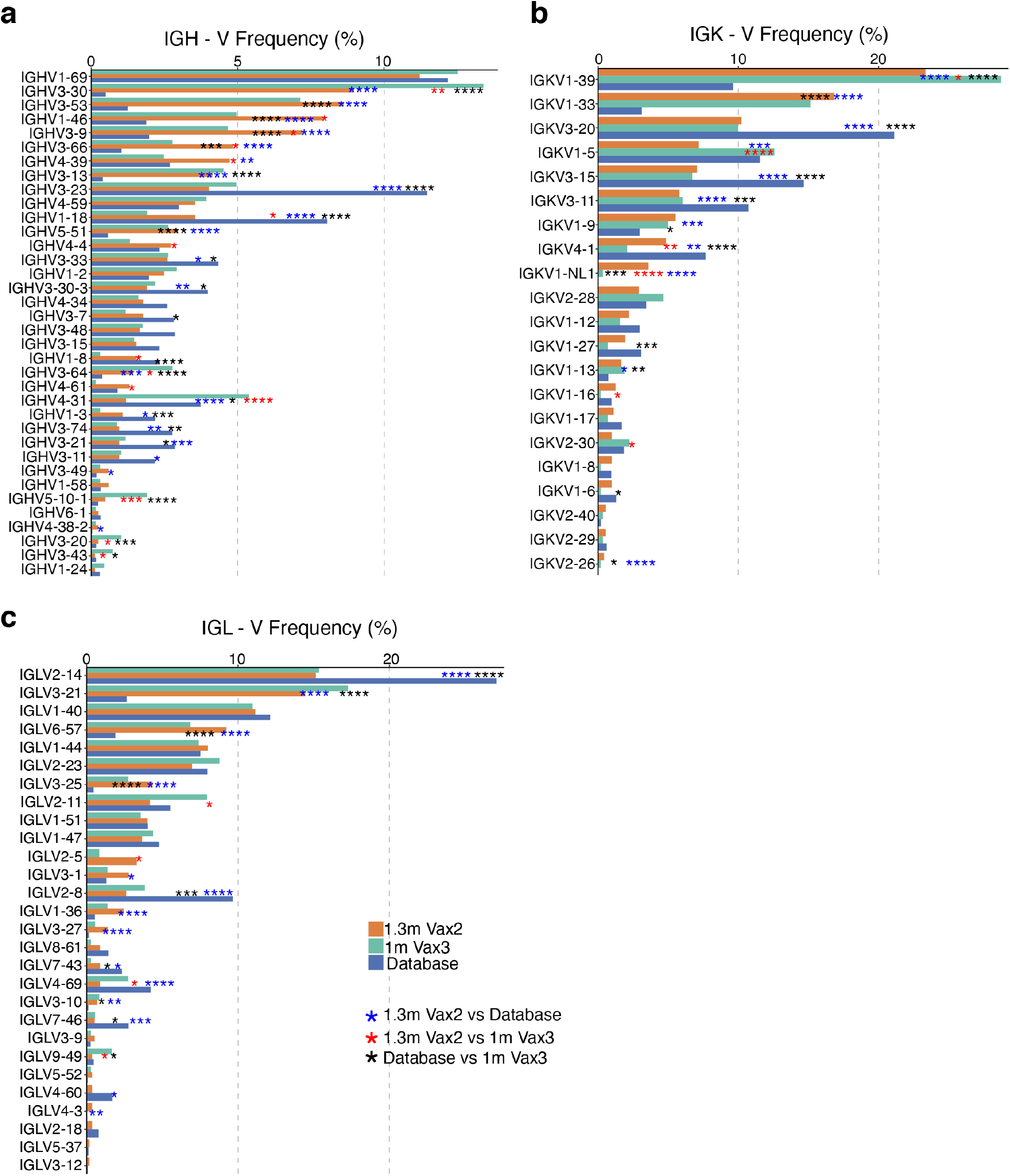
Frequency distribution of human V genes. **a-c** Comparison of the frequency distribution of human V genes for heavy chain and light chains of anti-RBD antibodies from this study and from a database of shared clonotypes of human B cell receptor generated by Cinque Soto et al^41^. Graph shows relative abundance of human IGVH (**a**), IGVK (**b**) and IGVL (**c**) genes Sequence Read Archive accession SRP010970 (blue), 1.3m-Vax 2 antibodies (orange), and 1m-Vax3 antibodies (green).

**Extended Data Fig. 4:**
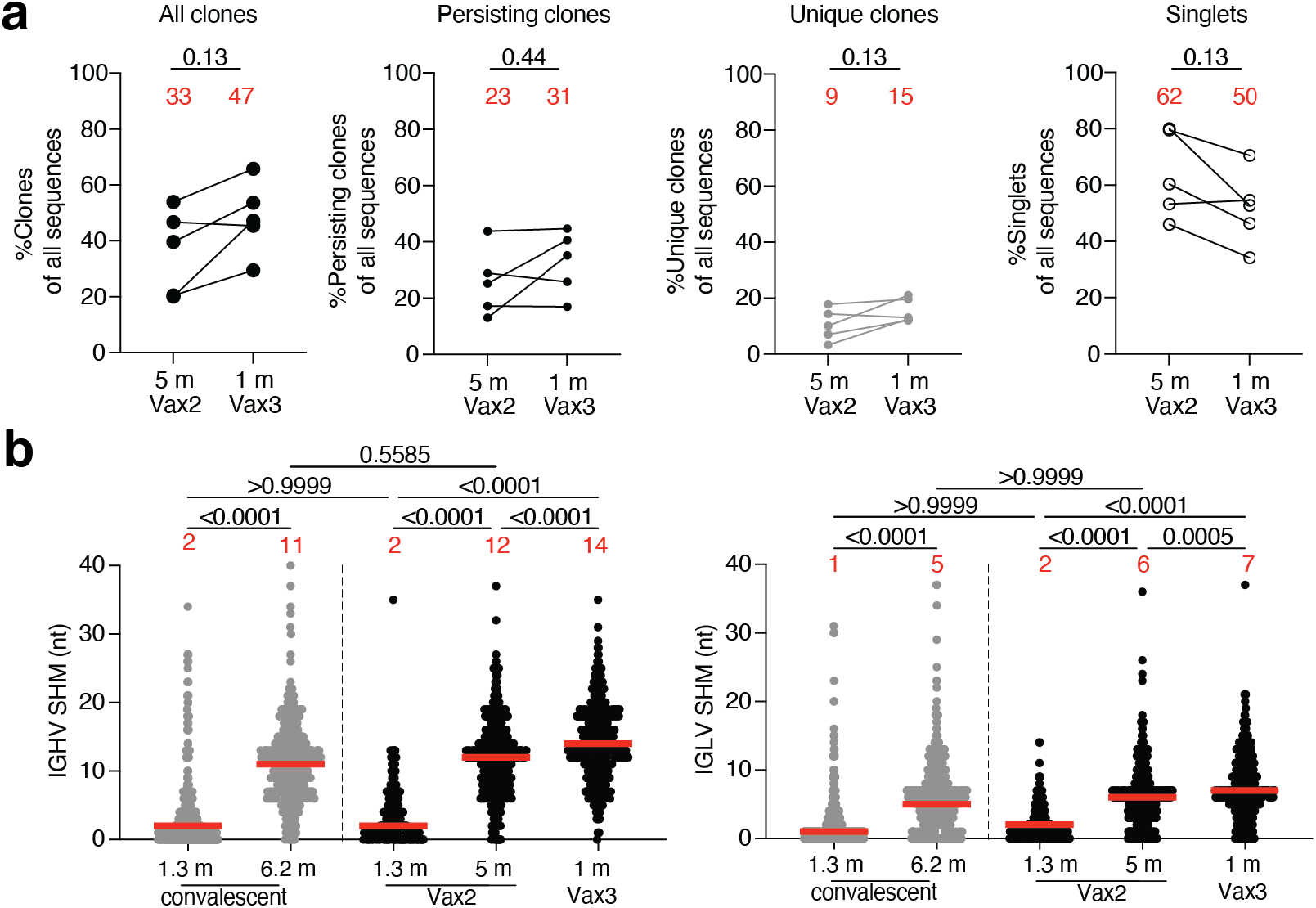
Clonality and somatic hypermutation of anti-SARS-CoV-2 RBD antibody clones after third vaccination booster. **a**, Graphs show relative fraction of clones, persisting clones, unique clones and singlets among all antibody sequences in n=5 individuals 5m after the 2^nd^ and 1 month after the 3^rd^ dose. **b**, Number of nucleotide somatic hypermutations (SHM) in the *IGHV* (left panel) and *IGLV* (right panel) in the antibodies illustrated in **Fig. 2c** for vaccinees after 1.3- and 5-months post-Vax2^9,10^ and 1 month after Vax3, compared to the number of mutations obtained after 1.3^18^ or 6.2^19^ months after infection (grey).

**Extended Data Fig. 5:**
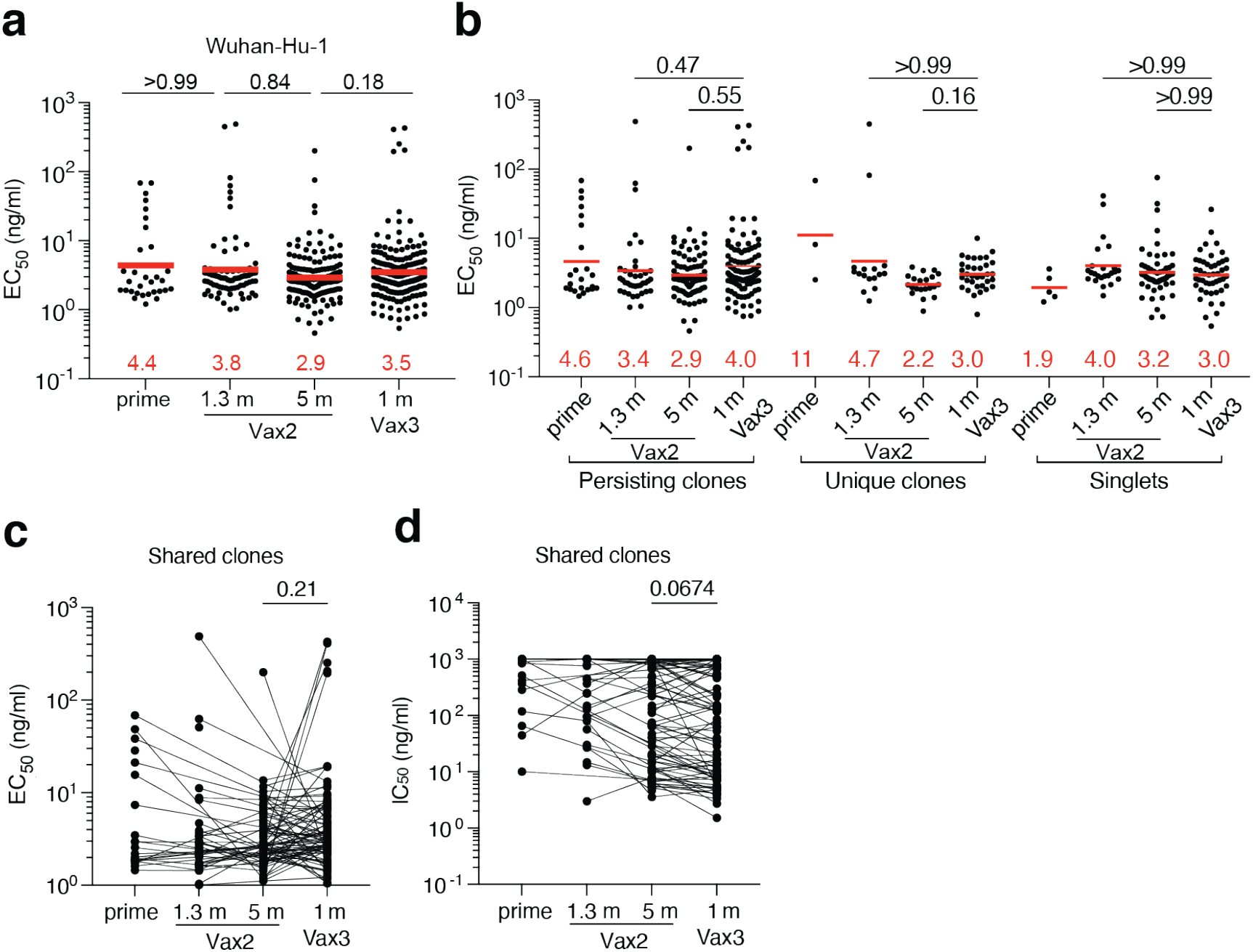
Anti-SARS-CoV-2 RBD monoclonal antibodies. **a**, Graphs show half-maximal concentration (EC_50_) of n=459 monoclonal antibodies measured by ELISA against Wuhan-Hu-1 RBD after prime^10^, 1.3- and 5-months post-Vax2^9,10^, and 1 month after Vax3. **b**, Graph showing EC50 of monoclonal antibodies as categorized as either persisting clones detected at multiple time points, unique clones where sequences were clonally expanded but detected at a single time point, or singlets were mAbs were derived from sequences detected once at a single time point. Graph showing **c**, EC_50_ of monoclonal antibodies or **d**, IC_50_ neutralizing activity from antibodies derived from shared clones only. Lines connect the related clones at the indicated time point. Red bars and numbers in **a**, and **b**, indicate geometric mean values. Statistical significance was determined by two-tailed Kruskal Wallis test with subsequent Dunn’s multiple comparisons. All experiments were performed at least twice.

**Extended Data Fig. 6.**
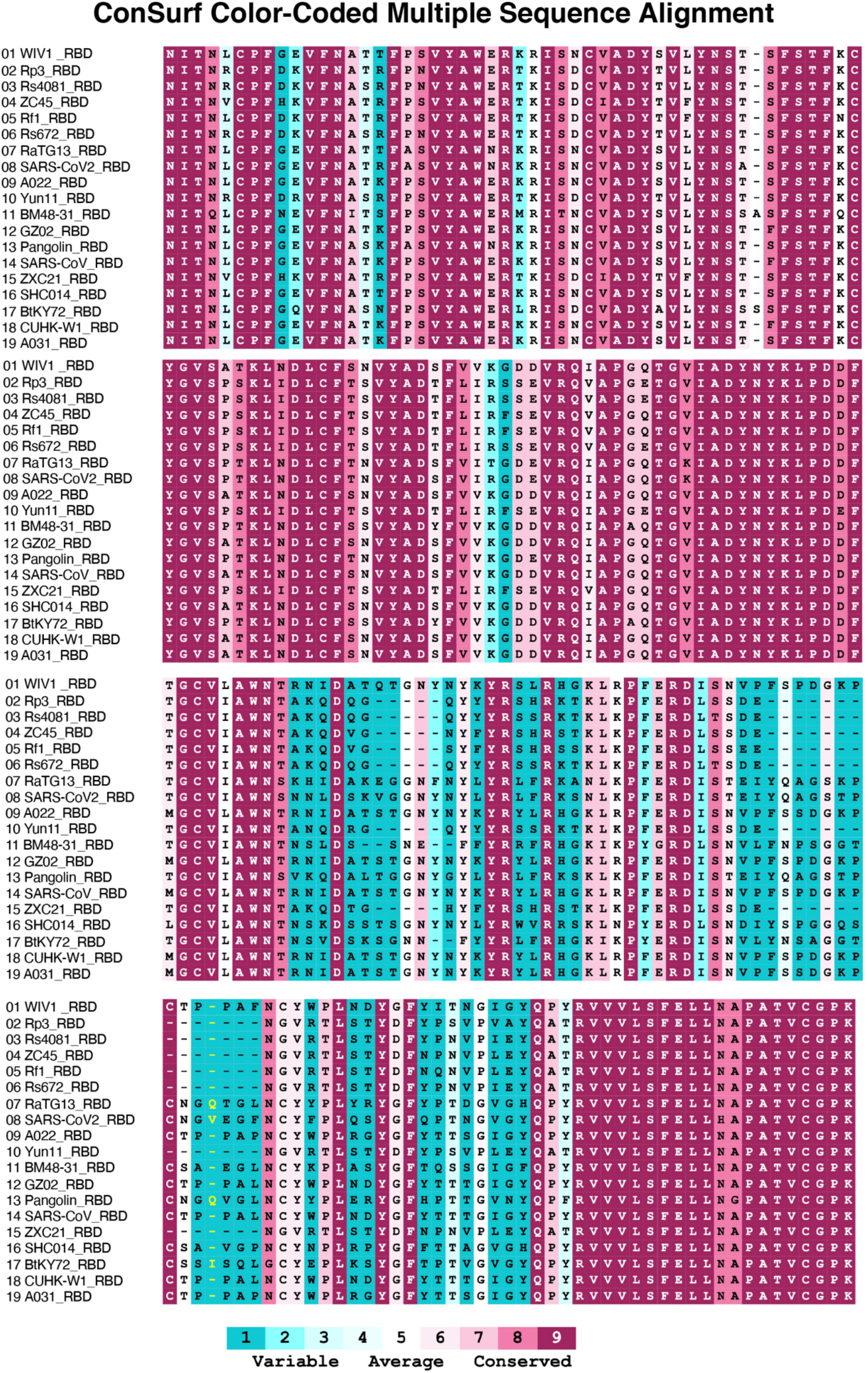
Multiple sequence alignment of RBDs. Sequences used for the alignment are the RBDs of WIV1(GenBank: KF367457.1), Rp3(UniprotKB:Q3I5J5), Rs4081(GenBank: KY417143.1), ZC45 (GenBank: AVP78031.1), Rf1(GenBank: DQ412042.1), Rs672(GenBank: ACU31032.1), RaTG13(GenBank: QHR63300.2), SARS-CoV2 (GenBank: MN985325.1), A022(GenBank: AAV91631.1), Yun11 (GenBank: JX993988.1), BM48-31(NCBI Reference Sequence: NC_014470.1), GZ02(GenBank: AAS00003.1), Pangolin(GenBank: QIA48632.1), SARS-CoV(UniProtKB:P59594), ZXC21(GenBank: AVP78042.1), SHC014(GenBank: KC881005.1), BtKY72(GenBank: KY352407.1), CUHK-W1(GenBank: AAP13567.1), and A031(GenBank: AAV97988.1). Multiple sequence alignment of RBDs was processed by Clustal Omega^43^. Sequence conservation was calculated by the ConSurf Database^44^.

## Supplementary Information

**Supplementary Table 1:** Individual participant characteristics.

**Supplementary Table 2:** Sequences of anti-SARS-CoV-2 RBD IgG antibodies.

**Supplementary Table 3:** Sequences, half-maximal effective concentrations (EC_50_s) and inhibitory concentrations (IC_50_s) of cloned monoclonal antibodies.

**Supplementary Table 4:** Binding and Neutralization activity of persisting clones

